# Increasing muscle speed drives changes in the neuromuscular transform of motor commands during postnatal development in songbirds

**DOI:** 10.1101/2020.02.19.955799

**Authors:** Iris Adam, Coen P.H. Elemans

## Abstract

Progressive changes in vocal behavior over the course of vocal imitation leaning are often attributed exclusively to developing neural circuits, but the effects of postnatal body changes remain unknown. In songbirds, the syrinx transforms song system motor commands into sound, and exhibits changes during song learning. Here we test the hypothesis that the transformation from motor commands to force trajectories by syringeal muscles functionally changes over vocal development in zebra finches. Our data collected in both sexes show that only in males, muscle speed significantly increases and that supralinear summation occurs and increases with muscle contraction speed. Furthermore, we show that previously reported sub-millisecond spike timing in the avian cortex can be resolved by superfast syringeal muscles and that the sensitivity to spike timing increases with speed. Because motor neuron and muscle properties are tightly linked, we make predictions on the boundaries of the yet unknown motor code that correspond well with cortical activity. Taken together, we show that syringeal muscles undergo essential transformations during song learning that drastically change how neural commands are translated into force profiles and thereby acoustic features. We propose that the song system motor code must compensate for these changes to achieve its acoustic targets. Our data thus supports the hypothesis that the neuromuscular transformation changes over vocal development and emphasizes the need for an embodied view of song motor learning.

**Significance statement:** Fine motor skill learning typically occurs in a postnatal period when the brain is learning to control a body that is changing dramatically due to growth and development. How the developing body influences motor code formation and vice versa remains largely unknown. Here we show that vocal muscles in songbirds undergo critical transformations during song learning that drastically change how neural commands are translated into force profiles and thereby acoustic features. We propose that the motor code must compensate for these changes to achieve its acoustic targets. Our data thus support the hypothesis that the neuromuscular transformation changes over vocal development and emphasizes the need for an embodied view of song motor learning.

## Introduction

Animal behavior results from system-wide interactions between nervous system, muscles, body, and surrounding environment (Chiel and Beer, 1997; Nishikawa et al., 2007). During motor skill learning the brain strives to generate activation patterns to reliably cause behaviors while the body exhibits large changes due to growth and training (Adam and Elemans, 2019). Because of the long extent (months to years) of fine motor skill learning little is known about how motor code changes and how the developing body contributes to behavioral changes. Recent work shows that small changes in biomechanics can cause behavioral state changes previously attributed to neural drive alone (Zhang et al., 2019). It is thus essential to include biomechanics when studying motor control (Nishikawa et al., 2007; Tytell et al., 2011) and especially over developmental timescales (Zhang and Ghazanfar, 2018).

An excellent model to study motor skill learning is vocal imitation leaning in songbirds. In zebra finches, males learn how to sing early in life by first memorizing a tutor song, which they match acoustically through trial- and-error learning over 60 days (Tchernichovski et al., 2001). Vocal output is controlled by well-characterized neural circuitry (aka the song system) that consists of two main pathways: i) the anterior forebrain pathway (AFP) necessary to learn and maintain song (Scharff and Nottebohm, 1991) and ii) the motor pathway, which encodes the learned vocalizations and is needed throughout life to produce them. In the avian cortex, the motor pathway produces millisecond-scale precisely-timed complex sequences of motor commands (Chi and Margoliash, 2001; Hahnloser et al., 2002) that result in force trajectories by respiratory and superfast vocal muscles (Elemans et al., 2008; Elemans, 2014). The transform from spikes to force is often referred to as the neuromuscular transform (NMT) (e.g. (Zhurov and Brezina, 2006)). During song, these force trajectories directly modulate pressure in the respiratory system (Srivastava et al., 2017), and the position and tension of vibrating structures in the sound-producing organ, the syrinx, thereby altering the vocal output (Riede and Goller, 2010; Elemans, 2014). Thus, via the NMT, motor commands of the song system directly drive the acoustic features of song.

The changes in song observed over the course of vocal imitation learning are typically attributed to changes in neural drive (Jarvis, 2004; Fitch et al., 2010). However, in this same period the syrinx and particularly the syringeal muscles that execute motor commands from the song system are also undergoing significant changes (Adam and Elemans, 2019). One change directly influencing the NMT is that force responses of syringeal muscles double in speed when stimulated by single spikes (Mead et al., 2017). The observed changes seem driven predominantly by use, suggesting that extensive training of syringeal muscles is essential to achieve their maximal speed (Adam and Elemans, 2019). However, in vivo muscle contractions underlying behavior are never elicited by single spikes, but by spike trains with variable inter-spike-intervals (ISIs), and it remains unknown if the observed changes in muscle speed over development functionally alter force profiles and thereby behavioral output.

Here we test the hypothesis that muscle properties relevant to the NMT change over vocal development in male songbirds. We investigated three critical NMT features related to activation by spike trains: 1) speed, 2) supralinearity, and 3) response to sub-millisecond precision spike timing. We show that all speed parameters of twitch and tetanic contractions significantly increase over vocal development. Furthermore, we show that supralinear summation occurs and increases over development. Lastly, we show that sub-millisecond spike timing can be resolved by syringeal muscles and that spike timing sensitivity increases with speed. Taken together, our data shows conclusively that the NMT changes over vocal development. We propose that song system must compensate for NMT changes to achieve correct song targets, such as pitch trajectories over syllables. Because motor neuron and muscle properties are tightly linked, our data allows for predictions on the boundaries of the yet unknown motor code that correspond well to cortical activity.

## Material and Methods

### Experimental Animals

All data presented here was collected in zebra finches (*Taeniopygia guttata*). Birds were raised in individual breeding cages together with their biological parents and siblings. In total 80 zebra finches were used in this study (m: 55, f: 25). All animals were kept on a 12:12h light:dark-cycle and provided with food and water ad libitum. All experiments and procedures were performed in accordance with the Danish Animal Experiments Inspectorate (Copenhagen, Denmark).

To quantify speed-related contraction parameters of syringeal muscles over the course of vocal development, we extracted muscle fiber bundles from *musculus tracheobronchialis dorsalis* (DTB) of male syrinxes at four time points: at 25 days post hatch (DPH), right before the start of the sensorimotor phase (Roper and Zann, 2006), at 50DPH during plastic song, at 75DPH, after the sensory phase has ended and at 100DPH when song is crystalized and the birds are considered adults (Immelmann, 1984). We included a cohort of age-matched females for comparison. Sex was determined by dissection post mortem (25DPH) or plumage (all other ages).

### In Vitro Muscle Physiology

Muscle fiber bundles were prepared and stimulated *in vitro* to record isometric force responses as previously described (Elemans et al., 2008; Srivastava et al., 2017). In brief, birds were sacrificed by an isoflurane overdose and the syrinx was exposed, isolated and pinned down on Sylgard-covered Petri dishes in cold, oxygenated Ringers solution. Fiber bundles were obtained randomly from the left or right DTB. Muscle preparations were mounted in a temperature-controlled bath, which was continuously supplied with oxygenated Ringers solution. The rostral end of the preparation was fixed to a force transducer (Model 400A, Aurora Scientific) and the caudal end to a micromanipulator, which was used to control length of the preparation. After mounting the muscle preparation in the bath chamber, it was allowed to rest for 20 minutes. Field stimulations were applied using platinum electrodes. Force and stimulation signals were low-pass filtered at 10 kHz (EF120 BNC through-feed low-pass filter, Thor Labs) and digitized at 20 or 40 kHz (NI DAQ Board PCI-MIO-16E4, National Instruments). All stimulations were carried out at 39.0±0.1°C. All software to control the setup and record data was written in Matlab (MathWorks, RRID:SCR_001622).

### Stimulation protocols

First, we optimized stimulation amplitude (pulse width of 300 μs) for maximal force production. Second, we applied 100 ms pulse trains ranging from 100 to 800 Hz in 100 Hz steps, to determine the force frequency curve and the stimulus frequency at which maximal tetanic force was produced. Lastly, muscle length was optimized for maximal force for each preparation.

A critical gap in our knowledge of the song system is that virtually no information is available about firing characteristics of the motor neurons during song. The output nucleus of both the AFP and motor pathway is the robust nucleus of the arcopallium (RA), which sends direct projections to the motor neurons located in the hypoglossal nucleus nXII (Nottebohm et al., 1976; Wild, 1993a, b). Motor neurons in the tracheosyringeal part of nXII (nXIIts) innervate the syringeal muscles (Wild, 1997). Although multiunit recordings of nXIIts have been reported (Williams and Nottebohm, 1985; Otchy et al., 2019), no firing characteristics of single units in nXIIts that comprise the actual motor code exist to our best knowledge. Thus, while neural coding in the song system upstream of the motor neurons is well described, virtually no information is available on the motor code controlling the vocal organ. High quality neural data has been obtained from single units in the motor cortex, i.e. nucleus RA, which is directly driving the syrinx motor neurons in nXIIts and we therefore used RA firing statistics to design our stimulation patterns. Over song development, RA firing activity develops from variable firing with a wide ISI distribution to sparse, bursty firing, that is precisely locked to song and has a very narrow ISI distribution (Leonardo and Fee, 2005; Olveczky et al., 2011).

For each muscle preparation we performed several stimulation experiments in series in the order described below. Each experiment consisted of several stimulation patterns (described in the section below for each experiment). Each iteration was started by a tetanic pulse train (100 for experiment 1 and 50 ms for other experiments) at optimal stimulus and length followed by 3 minutes rest. Subsequently, all stimulation patterns of one experiment were applied in random order with two minutes rest in-between. Each experiment was then repeated for several iterations (specified below for each experiment) and patterns were re-randomized within each iteration.

Experiment 1: Isometric stress and muscle speed. Each iteration consisted of a 100 ms stimulation at the frequency producing maximal force followed by 4 single twitches. The experiment was run for at least two iterations.

Experiment 2: Supralinear force summation. We used 4 stimulation patterns consisting of two spike patterns with ISIs of 2, 4, 6 and 8 ms, covering the range of ISIs most commonly observed in RA (Olveczky et al., 2011). The experiment was run for at least three iterations.

Experiment 3: Spike timing. To investigate the effect of spike timing on force responses, we designed a three-spike stimulation paradigm (based on(Srivastava et al., 2017)) consisting of 29 stimulation patterns based on four different frequencies (200, 400, 600, 800 Hz). To introduce spike timing changes without changing the overall rate, the middle spike was moved in small increments (range 0.1 – 1 ms) from its centered position towards the first or last spike. The stimulation patterns by frequency were as follows: At 200 Hz we used 11 stimulation patterns with ISIs between the first and second spike (ISI1) of 0.4, 0.5, 1, 2, 3, 4, 5, 6, 7, 8, 9 ms. At 400 Hz we used 7 stimulation patterns with ISI1s of 0.4, 0.5, 1, 2, 2.5, 3, 4 ms. At 600 ms we used 6 stimulation patterns with ISI1s of 0.4, 0.5, 1, 1.66, 2, 3 ms. At 800 ms we used 5 stimulation patterns with ISI1s of 0.4, 0.5, 1, 1.25, 2 ms. The experiment was run for at least 2 iteration.

Experiment 4: Refractory period. Stimulation patterns consisted of 8 two spike patterns with ISIs ranging from 0.3 - 1.5 ms in 0.2 ms steps and a single pulse stimulation as control. The experiment was run for at least 2 iterations. Data was acquired on 13 additional animals and acquired directly after experiment 1.

### Data analysis

All data analysis was carried out in R (R Project for Statistical Computing, RRID:SCR_001905).

#### Accounting for force decay

The above described experiments amounted to between 16 to 37 individual stimulation patterns and acquiring all data typically took more than 12h, during which the force output of the preparation decreased due to dying muscle fibers. To correct for this force decrease, we normalized all force data from our stimulation patterns within one experiment. For normalization we used the linear regression between the maximal force of the 50 ms tetanic contractions, which were part of each iteration, and experimental time.

#### Isometric stress

Stress was calculated as F/A_CSA_ of the muscle, where the cross-sectional area A_CSA_ was estimated from the resting length L_0_, the dry weight (dry-wet conversion factor: 5) and density (1060 kg/m^3^ from (Mendez and Keys, 1960)) of the muscle fibers.

#### Muscle speed, single stimulations

The full width at half maximal (FWHM) force of single twitches was defined as the time from crossing 50% force increase to decrease, and is the same parameter as previously reported t_50-50_ (Elemans et al., 2004; Srivastava et al., 2017). The time to maximal force (F_max_) was defined as the time from stimulation to reaching the peak of the twitch. Latency was defined as the time from stimulation to the force signal crossing the mean plus 5 times S.D. of a 50 ms segment of the force prior to stimulation. All values reported are the average of 7 twitch contractions. FWHM of another syrinx muscle (VTB) was extracted in 12 additional animals from previously published data (Mead et al., 2017).

#### Muscle speed, tetanic contractions

We fitted a mathematical model (Watanabe et al., 2010) to the Fusion Index-frequency-curve (FFC) and force-frequency-relation (FFR) using the *n/s* function in R to extract the frequencies corresponding to 10% fusion (start of summed responses), 90% fusion (*f_fused_*) and 90% of maximal tetanic force (*f_Tet_*).

#### Supralinearity

We normalized the maximal force amplitude of all force traces to the F_max_ of the first peak of the 8 ms pattern within each iteration. A twitch model (Raikova and Aladjov, 2002) was fitted and used to calculate linear summations by adding two modeled twitches offset by the ISIs used for stimulation. Supralinearity was quantified as the ratio between the normalized and modelled F_max_ as well as the ratio of the area under the curve (AUC) of the normalized force data and the summation model. The AUC was calculated using the *AUC* function from the *DescTools* package (Signorell and al., 2019).

#### Spike timing

To quantify the variability introduced by changing spike timing, the coefficient of variation was calculated for the timing (CV_Timing_) as well as for the amplitude (CV_Amplitude_) of F_max_ per stimulation frequency and animal. All iterations where force decayed by more than 20% compared to the start of the experiment were excluded from analysis.

#### Estimation of pitch changes

The fundamental frequency of sound is often called “pitch”. Even though pitch is a perceptual measure, that is not always equivalent to fundamental frequency, we will refer to the fundamental frequency as pitch to avoid confusion between stimulation and sound frequencies. To illustrate the effect of spike timing manipulations on sound production, we estimated the pitch changes which would occur if the entire *musculus syringealis ventralis* (VS) would be stimulated simultaneously with the three-pulse patterns during sound production. We focused on VS, as it is well established that VS controls pitch (Goller and Suthers, 1996; Elemans et al., 2015; Srivastava et al., 2015), while the function of DTB is less clear. VS force was calculated for each animal by dividing the stress data by the CSA of VS (1,05 mm^2^) (Adam et al., 2020). Pitch was calculated from force data using the force - pitch transformation (pitch = 26.6\**F*_VS_) of the VS muscle (Adam et al., 2020). The average pitch difference per animal was calculated as the difference between the minimal and maximal pitch values per bird across all stimulation patterns of the spike timing protocol.

#### Refractory period

F_max_ values were extracted from all force responses and normalized to F_max_ of the control twitch within each iteration. To estimate the refractory period (t_Ref_), we fitted a log-logistic model to the normalized F_max_ values using the *drm* function of the *drc* package (Ritz et al., 2015):

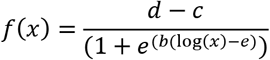

The t_Ref_ was estimated as the ISI, where the normalized maximal force was 10% higher than of a single twitch response using linear interpolation (Boerio et al., 2005).

### Experimental design and statistical analyses

All data is reported as average ± standard deviation; N refers to individual animals. Statistical significance was accepted for *p* < 0.05.

#### Statistical tests

Linear mixed models with subsequent Tukey’s posthoc comparison were used to assess statistical difference in the FWHM (Fig. 1B), F_max_ (Fig. 2B) and supralinearity (Fig. 2E) data sets. Statistical design for each data set can be found in the respective figure captions. ANOVA with subsequent Tukey’s posthoc test was used to detect significant differences between age groups in the *f_fused_* (Fig. 1F), *f_max_* (Fig. 1G), time to F_max_ (Fig. 1C) and latency (Fig. 1D) data sets and between stimulation frequencies and CV_Amplitude_ and CV_Timing_ (Fig. 3B). Linear models were employed to test the correlation with *f_fused_* for FWHM (Fig. 1H), supralinearity (Fig. 2F+G) and CV_Amplitude_ and CV_Timing_ (Fig. 3C). Welch t-test was used to test for sex differences in t_Ref_ (Fig. 4C).

**Figure 1:**
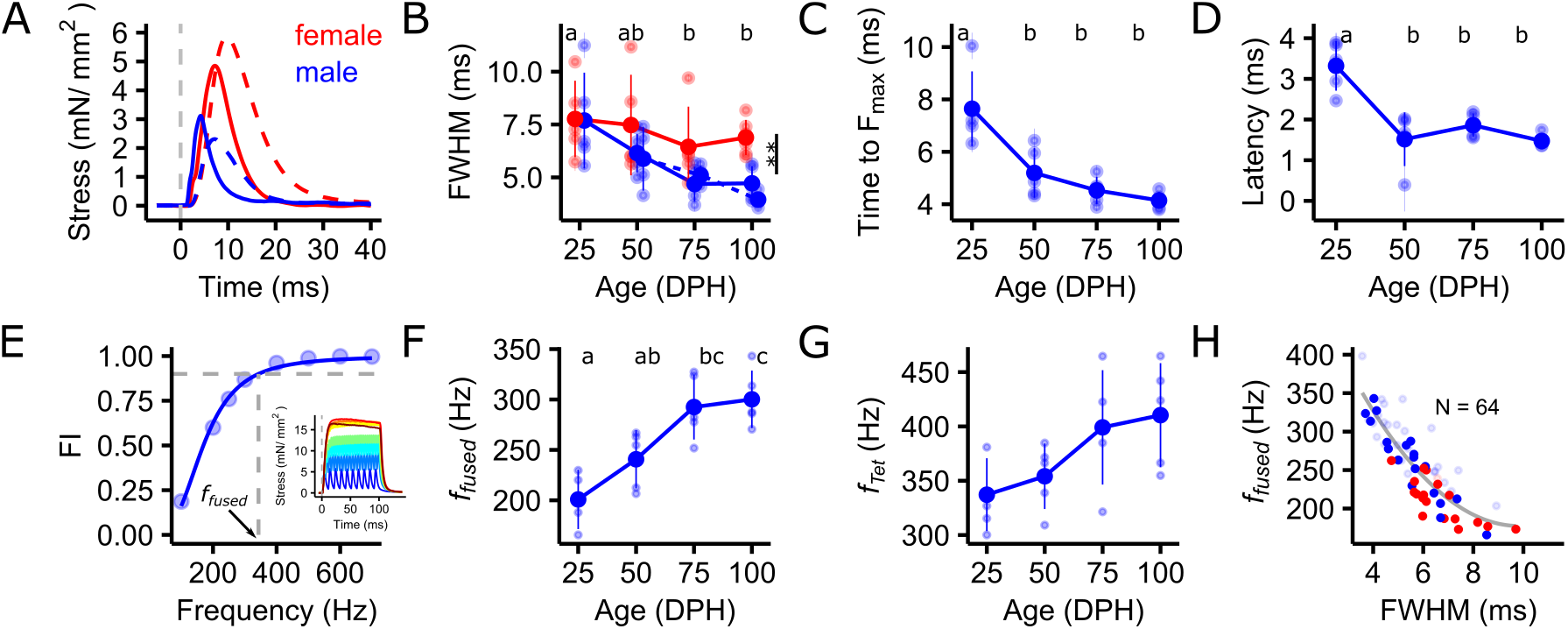
Contraction speed increases over development in male songbirds. *A*, Twitch contraction speed was higher in adult males compared to juveniles or adult females as illustrated by the narrower force peak. Examples of twitch responses of juvenile (dashed lines) and adult (solid lines) male (blue) and female (red) zebra finch syrinx muscles. The dashed, vertical, grey line indicates the time point of stimulation. *B*, Full width at half maximal (FWHM) force decreased over vocal development in males, but not in females (LMM: FWHM~sex + age + (1|animal), sex: df(1), F=7.5, p<0.01, age: df(3), F=3.9, p<0.05, animal: df(1), LRT=835, p<0.001). *C*, Time to F_max_ decreased significantly between 25DPH and 50DPH (ANOVA: F=14.98, df(3,16), RSE=0.91, p <0.001), as did the latency from stimulation to the force onset (D) (ANOVA: F=16.5, df(3,16), RSE=0.48, p <0.001). *E*, A mathematical model was fitted to the Fusion Index frequency curve (FFC) and the force-frequency-relation (FFR) to extract the frequency when responses start to fuse (FI=0.1), reach 90% fusion (*f_fused_*) and when force reaches 90% of the maximum force (*f_Tet_*), respectively. Depicted is an example of an FFC of an adult male. Crossing of the grey dashed line with the x-axis indicates *f_fused_*. The inset depicts the force responses to stimulation with spike trains from 100 to 700 Hz (encoded by color) used to calculate FI. *F, f_fused_*) and (G) *f_tet_* both increased significantly over vocal development. (*f_fused_*: ANOVA: F=10.91, df(3,15), RSE=29.84, p <0.001; *f_Tet_*: ANOVA: F=3.48, df(3,16), RSE=42.14, p=0.041). *H*, FWHM and *f_fused_* are highly correlated (N=64, LM: *f_fused_* ~ FWHM, p< 0.001, RSE=51.66, df(63)). Solid points indicate data from preparations depicted in *B*, *C*, *D*, *F*, *G*, while transparent points are derived from additional preparations of other syringeal muscles, partially published in (Mead et al., 2017). Unshared lowercase letters above plots indicate post-hoc differences with significance level p<0.05. A horizontal jitter was applied to data points in *B* for clarity.

**Figure 2:**
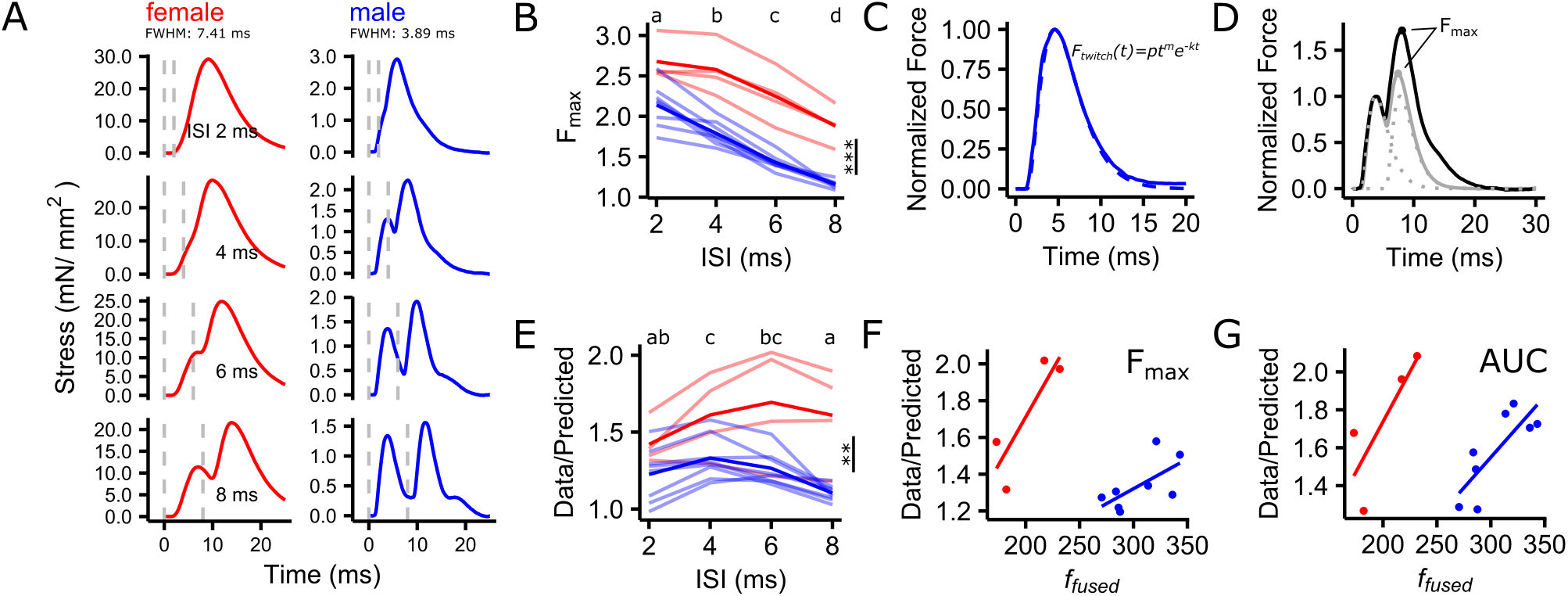
Force summation is supralinear and depends on contraction speed. *A*, The shape of force trajectories changed dramatically with muscle contraction speed (male vs female) when driven by identical stimulation patterns. This is illustrated by examples of force responses of a female (red) and a male (blue) syrinx muscle to stimulations with two pulses with ISIs of 2, 4, 6 and 8 ms. The dashed, vertical, grey lines indicate the time points of stimulation. *B*, Maximal force relative to a single twitch decreased with increasing ISI and was higher in males compared to females (LMM: Normalized F_max_ ~ sex + ISI + (1 | animal), sex: df(1), F=45.4, p<0.001, ISI: df(3), F=102.3, p<0.001, animal: df(1), LRT=16.3, p<0.001). Transparent lines represent individual preparations, solid lines represent means. *C*, Twitch responses were fitted to a phenomenological model (Raikova and Aladjov, 2002). *D*, Recorded force responses (black) were compared to the summation model (grey) derived by summing two twitch models (dashed line). The amount of supralinearity was defined as the ratio between F_max_ or the AUC of the measured data and the summation model. *E*, The amount of supralinearity depended on ISI and was higher in females compared to males (LMM: Supralinearity~ sex + ISI + (1 | animal), sex: df(1), F=11.5, p<0.01, ISI: df(3), F=5.2, p<0.01, animal: df(1), LRT=21, p<0.001). On average, the maximum occured at a lower ISIs in males (4 ms) compared to females (6 ms). (*F*, *G*), Maximal supralinearity increased with *f_fused_*, for both F_max_ (LM: Ratio F_max_~ *f_fused_* + sex, *f_fused_*: df(2), t=3, p<0.01, sex: df(9), t=-4.3, p<0.01) and AUC (LM: Ratio AUC~ *f_fused_* + sex, *f_fused_*: df(2), t=3.8, p<0.01, sex: df(9), t=-4.1, p<0.01). Unshared lowercase letters above plots indicate post-hoc differences with significance level p<0.05.

**Figure 3:**
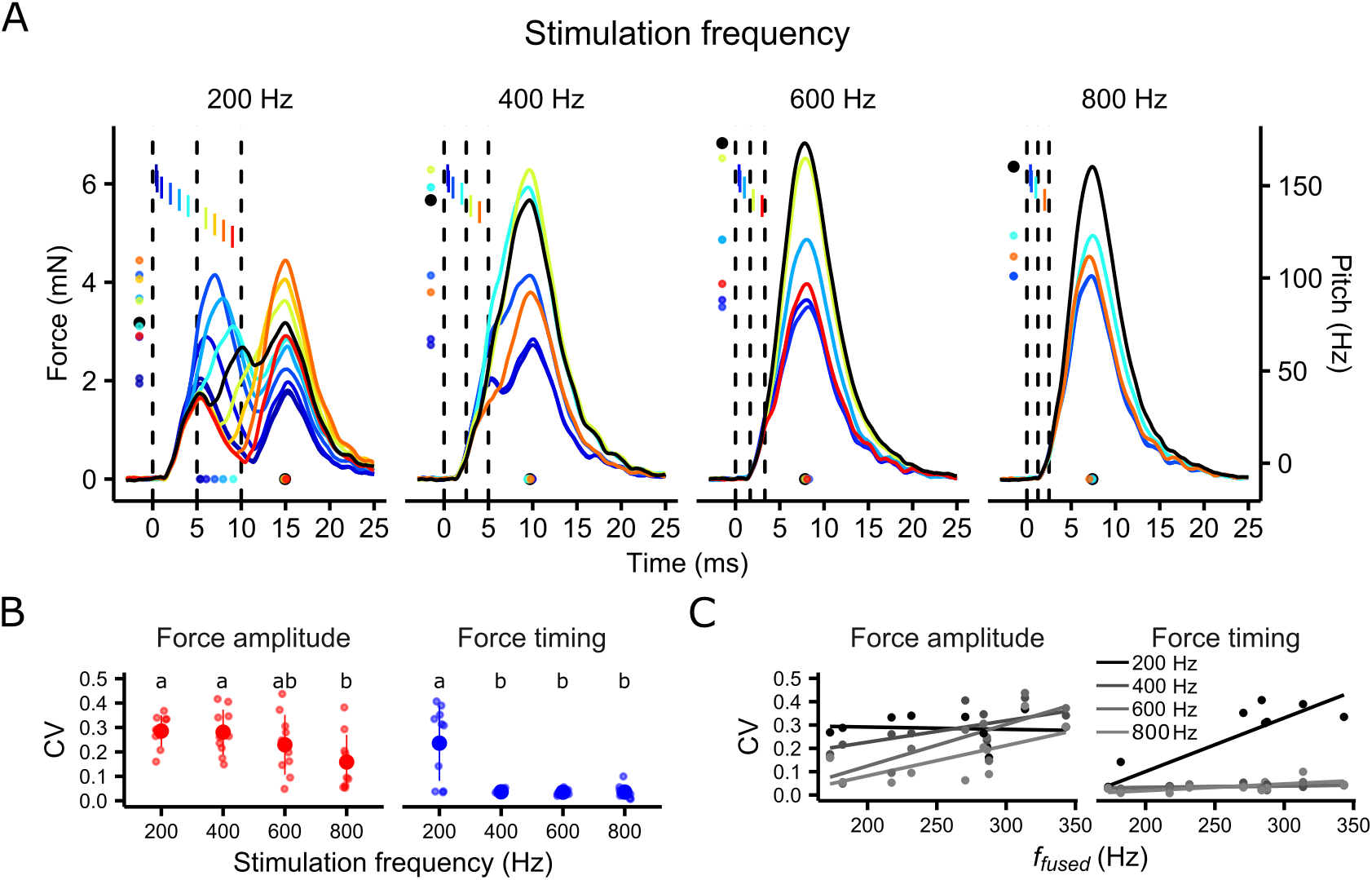
Sub-millisecond changes in spike timing affect magnitude and timing of maximal force development in superfast syrinx muscles. *A*, Three-spike patterns with timing changes to the middle spike cause changes in both timing and amplitude of force profiles as illustrated by example raw data of a male syrinx muscle (FWHM: 4.45 ms, *f_fused_*: 284 Hz). Lines depict the average normalized force response over 3 renditions. Indicated are the timing (dots at bottom) and amplitude (left) of F_max_. Vertical dashed bars indicate the timepoints of stimulation with regular ISI and the colored bars indicate the stimulation time point of the middle spike when offset from the centered position. Color encodes the relative offset of the middle spike from its regular position. *B*, The variability introduced by changes to spike timing depended on the base stimulation frequency and was generally higher at 200 Hz (CV_Amplitude_: 0.29 ± 0.06, CV_Timing_: 0.24 ± 0.15) compared to 800 Hz (CV_Amplitude_: 0.16 ± 0.11, CV_Timing_: 0.04 ± 0.03) (CV_Amplitude_: ANOVA: F=3.47, df(3,36), RSE=0.100, p <0.05, CV_Timing_: ANOVA: F=16.14, df(3,36), RSE=0.079, p <0.001). *C*, CV_Amplitude_ significantly depended on muscle speed (*f_fused_*) at base stimulation frequencies of 600 and 800 Hz (600 Hz: N=10, LM: CV_Amplitude_ ~ *f_fused_*, p=0.004, F=15.41, 800 Hz: N=10, LM: CV_Amplitude_ ~ *f_fused_*, p=0.04, F=5.96), while CV_Timing_ did so at 200 Hz (N=10, LM: CV_Timing_ ~ *f_fused_*, p=0.003, F=17.92). A horizontal jitter was applied to data points in *B* for clarity.

**Figure 4:**
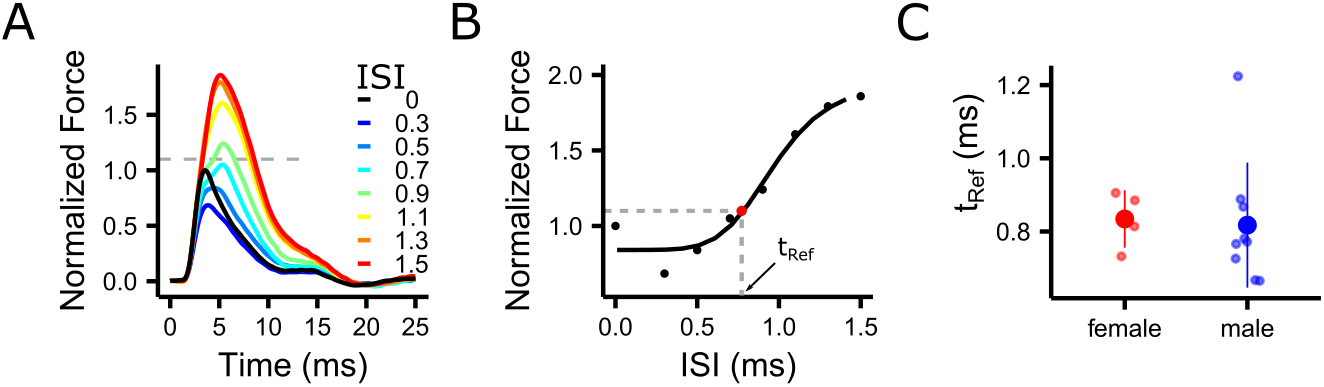
The refractory period of syringeal muscle fibers is sub-millisecond. *A*, Force summation decreases with ISIs below 1 ms and eventually disappears completely as shown by example force traces of stimulations with short ISIs of a male syrinx muscle. Color encodes the ISI and the dashed grey line indicates the threshold used to estimate t_Ref_ *B*, t_Ref_ was calculated by fitting a log-logistic model and was defined as the ISI between two spikes, where the F_max_ is 10 % higher than F_max_ of a single twitch. C, t_Ref_ of male and female DTB was not significantly different between males and females (Welch t-test: t=0.242, df=10.87, p=0.81).

As the FWHM did not change significantly with developmental age in females, we did not analyze the female data for all further measures of muscle speed in the developmental data set. To increase the range of muscle speed in our data set quantifying supralinearity and the influence of spike timing, we included data from female DTB preparations.

All statistical comparisons were carried out in R (R Project for Statistical Computing, RRID:SCR_001905).

## Results

### Contraction speed of syringeal muscles increases over vocal development in male songbirds

We measured isometric contraction speed of vocal muscles over song learning using two commonly applied stimulation paradigms: i) single stimulations yielding a single twitch contraction and ii) stimulations with equidistant trains of spikes with varying frequencies (100 - 800 Hz) resulting in summed force responses (Fig. 1). At 25DPH, FWHM of single twitches did not differ significantly between males and females and was 7.7 ± 2.2 ms (N=5) and 7.8 ± 1.8 ms (N=5) respectively. Over development, the speed of female DTB muscles did not change significantly and FWHM values stayed similar until 100DPH (6.9 ± 0.8 ms, N=6) (Fig. 1*B*). However, in strong contrast, male DTB muscles became steadily faster and finally doubled in speed with force FWHM almost halved to 4.7 ±0.8 ms (N=5) at 100DPH, corroborating previous findings in another syringeal muscle (Mead et al., 2017). The time from stimulation to maximal force (F_max_) (Fig. 1*C*) and latency from stimulation to force onset (Fig 1*D*) also steadily increased over development but were already significantly different between 25DPH and 50DPH.

We measured three other muscle speed parameters by applying multiple equidistant stimulation trains: the stimulation frequency at which the force response i) starts to summate, ii) is fully fused (*f_fused_*) and iii) reaches maximal tetanic force (*f_Tet_*) (See Methods). The frequency at which twitches start to summate increased from 65.43 ± 6.24 Hz at 25DPH (N=4) to 75.26 ± 11.24 Hz in adult animals (N=5). *f_fused_* increased steadily and significantly from 25DPH (201 ± 29 Hz, N=4) to 100 DPH, reaching its maximum at 300 ± 29 Hz (N=5) (Fig. 1*F*). Likewise, *f_Tet_* increased from 337 ± 34 Hz at 25DPH (N=5) to 410 ± 48 Hz at 100DPH (N=5) (Fig. 1*G*). The *f_Tet_* was higher than *f_fused_* in all groups (seen by higher values in Fig. 1*F* compared to Fig. 1*G*). This means that force still increased as a function of stimulation frequency even after the force responses were fully fused. This has also been reported in other motor systems (Kernell, 1995; Orizio et al., 2004). Interestingly, in our large dataset of 64 animals we found that force FWHM accurately predicted the stimulation frequency at which fused tetanic contractions occurred, independently of age, sex and muscle (Fig. 1*H*). Taken together, all isometric twitch and tetanic speed parameters steadily and significantly increased over vocal development in male songbirds.

### Supralinear force summation increases with muscle speed

Activating muscle fibers with multiple spikes at sufficiently small ISIs leads to summed force responses that are larger than expected from simple algebraic summation of single twitches (Kernell, 1995). This intrinsic property of the NMT is often referred to as supralinear or nonlinear force summation (Kernell, 1995; Binder et al., 2010; Srivastava et al., 2017). In the songbird respiratory system, millisecond changes in spike timing can cause significant behavioral changes, that reflect and potentially even exploit these nonlinear properties of muscles (Srivastava et al., 2017). The same could be true in syrinx muscles, but we don’t know whether they also exhibit supralinear summation to begin with and if its magnitude changes depending on muscle speed. Therefore, we quantified the magnitude of supralinear force summation in syringeal muscles (See Methods).

First, when driven by identical stimulation patterns the shape of force trajectories changed dramatically with muscle contraction speed (Fig. 2*A*). Maximal force (F_max_) decreased significantly with increasing ISI (Fig. 2*B*)in all preparations and was significantly higher in females than in males for all ISIs tested (f: 2.3 ± 0.4, N=4; m: 1.6 ± 0.4, N=8). Second, force summation was supralinear in all preparations, as the observed force response had a higher F_max_ than expected from a summation model (Fig. 2*E*, See Methods). The magnitude of supralinear summation significantly depended on spike timing (Fig. 2*E*) corroborating previous findings in songbird respiratory muscles (Srivastava et al., 2017). Third, the maximal magnitude of supralinear summation increased with muscle speed, irrespective of whether it was calculated from F_max_ (Fig. 2*F*) or AUC (Fig. 2*G*). Thus, taken together, the maximal magnitude of supralinear summation increases with speed and can thus be also expected to increase over development.

### Sub-millisecond changes in spike timing affect magnitude and timing of maximal force development

Timing of spikes relative to a behavior usually exhibits variability (Stein et al., 2005; Lisberger and Medina, 2015) and it has recently become apparent that the exact timing carries information (Stein et al., 2005; Sober et al., 2018). In songbirds, variability in RA indeed correlates to acoustic variability down to the millisecond scale (Sober et al., 2008; Tang et al., 2014). However, in RA firing precision is even higher and reported as 0.2 ms (Chi and Margoliash, 2001), but it is unknown whether sub-millisecond precision spike timing can pass through the syringeal muscle NMT and cause changes in force trajectories.

To address this question, we performed muscle stimulations with three-spike patterns introducing timing changes to the middle spike in steps from 1 to 0.1 ms (See Methods). These timing manipulations evoked changes in both the timing and amplitude of force profiles and the effect differed between stimulation frequencies (Fig. 3*A*). To quantify the amount of variability introduced by the timing manipulations, we calculated the coefficient of variation for the timing (CV_Timing_) and amplitude (CV_Amplitude_) of F_max_. Both, CV_Timing_ and CV_Amplitude_ were significantly higher at a stimulation frequency of 200 Hz compared to 800 Hz (Fig. 3*B*, *C*). While 200 Hz stimulation patterns caused variability in both timing and magnitude of F_max_, higher stimulation frequencies almost exclusively affected F_max_ amplitude (Fig. 3*B*), but not its timing (Fig. 3*C*).

Because the variability of force timing and amplitude introduced by changes in spike timing showed a wide variation between animals, we hypothesized that the effects could depend on muscle speed. To assess this hypothesis, we tested for the dependency of CV_Timing_ and CV_Amplitude_ on *f_fused_*, representing the intrinsic contraction speed of each animal. Overall, CV_Amplitude_ and CV_Timing_ increased significantly with *f_fused_* (Fig. 3*B*). However, as the stimulation frequency clearly influenced the magnitude of the CVs as well as the dependency on *f_fused_*, we calculated linear models for each stimulation frequency separately. CV_Amplitude_ significantly depended on *f_fused_* at 600 and 800 Hz and CV_Timing_ at 200 Hz (Fig 3*C*). These results mean that the quality of the effect of changes in spike timing are set by the relationship between stimulation frequency and muscle speed (i.e. *f_fused_*): Force timing is affected at stimulation frequencies below the *f_fused_* of adult male muscles (range: 173 – 343 Hz, median: 277 Hz), while force amplitude is affected at all stimulation frequencies. The quality of the effect on the other hand is set by muscle speed confirming our hypothesis, that the variability introduced by spike timing changes depends on the contraction speed of the muscle. Taken together, our results show that sub-millisecond spike timing affects the shape of force profiles. Syringeal muscles can thus resolve spike timing in the sub-millisecond range. Additionally, muscle speed sets the sensitivity of the muscle to changes in force timing as illustrated by the increase in variability with increasing muscle speed.

Even though spike timing changes in the sub-millisecond range affected F_max_ of force profiles, it is not given that these force changes result in changes in sound production within the physiologically relevant range. For illustration, we estimated the maximal pitch difference caused by stimulating VS with our three pulse patterns (Fig. 3*A*, See Methods) based on *in vitro* measurements of the VS force to pitch transform (Adam et al., 2020). We estimated an average force difference of 8.48 ± 5.84 mN (N=10) between the minimal and maximal F_max_ across all stimulation patterns, which translated to an estimated pitch difference of 225 ± 155 Hz well within the range detectable by zebra finches. As zebra finch females already detect changes in pitch below 1Hz (Lohr and Dooling, 1998), we conclude that spike timing changes in the range of sub-milliseconds might have behavioral correlates.

Changing spike timing in the high-frequency stimulation patterns (600 and 800 Hz) always resulted in a reduction of maximal force compared to the regular stimulation pattern (Fig 3*A*). We hypothesized that this decrease was caused by moving spikes into the refractory period of the muscle (t_Ref_), which is 2 - 5 ms for most skeletal muscles (Farmer et al., 1960). To estimate the unknown t_Ref_ of syringeal muscles we stimulated with two pulses with very short ISIs (Fig. 4*A*, See Methods). The t_Ref_ was extremely short with 0.83 ± 0.08 ms for female and 0.82 ± 0.17 ms for male syrinx muscles (female: N=4, male: N=9) and 3-7 times faster than regular skeletal muscle. Thus, despite the extremely short t_Ref_, we confirmed our hypothesis that reduction of maximal force due to spike timing changes at 600 and 800 Hz is caused by moving spikes into the refractory period of the muscle.

## Discussion

We show that the songbird syrinx undergoes essential transformations during song learning that drastically change how neural commands are translated into force profiles and thereby acoustic features. All isometric contraction speed-related features of male syrinx muscles, such as FWHM, fusion frequency (*f_fused_*) and *f_Tet_*, significantly increased from juveniles to adults. Interestingly, also the latency from stimulus to force production significantly decreased. Furthermore, we show that two additional NMT features change significantly with muscle speed. First, we establish that supralinear summation of twitch force occurs in syrinx muscles at ISIs in the millisecond range and that its magnitude increases with muscles speed. Interestingly, the magnitude is much lower in the faster male muscles compared to female muscles even at comparable muscle speeds, which indicates that additional factors affect supralinear force summation. Second, we show that the observed spike timing in the sub-millisecond range in cortical area RA (Chi and Margoliash, 2001) can be resolved by syringeal muscles. Force amplitude was affected at all spike rates tested, while changes in peak force timing were evoked only when the average spike rate was below *f_fused_*. Furthermore, the sensitivity to changes in spike timing was set by muscle speed. Taken together, our data thus provide strong support for our hypothesis that muscle properties relevant to the NMT change over vocal development in songbirds. The observed NMT changes are driven by syringeal muscle speed as it increases over vocal development. This implies that the extreme speed of the adult muscle is not available to juveniles during song learning.

The song template is assumed to be fixed during song learning (Mooney, 2009) and maintained through adulthood in closed-ended song learners (Brainard and Doupe, 2013). Individuals are trying to minimize perceptual error in trial-to-trial renditions (Fee and Scharff, 2010). The increasing syringeal muscle speed over development has the consequence that identical motor commands result in distinct force trajectories depending on postnatal age and thus alter behavioral output. Thus, if an individual bird aims to produce the same force trajectory during song development, we propose it consequently must dynamically alter its motor code to correct for NMT changes when trying to sing target syllables.

Even though the physiology of the song system neurocircuitry is well studied, the actual motor code, i.e. the firing patterns of single units in nXIIts, during singing are unknown (Williams and Nottebohm, 1985; Otchy et al., 2019). However, in many motor systems the electrical intrinsic properties of motor neurons and the biomechanical properties of the innervated muscle fibers are highly tuned (Zhurov and Brezina, 2006; Manuel et al., 2019). For example, the discharge rates of motor neurons typically fall within the range frequencies required to generate summed force responses by the muscle fibers. This is reflected by matching between the firing characteristics of the motor neurons and its muscle fiber force-frequency relationship (FFR) so that the observed range of firing rates modulates forces along the steepest portion of the FFR (Kernell, 1995). This phenomenon is called speed matching. Reversely, the speed properties of muscles thus set boundaries within which the motor code is operating. The intrinsic FFR property of a muscle thus allows for to predict the firing rates of the motor neurons innervating it.

Using our characterization of the biomechanical properties of syringeal muscles, we can make three predictions for the nXIIts motor neuron firing frequencies during singing in adult males. First, force summation begins at ISIs above 13.3 ms (i.e. 75 Hz). Thus, in order to activate syrinx muscles to produce summed force responses, firing rates have to be higher than 75 Hz. Second, spikes arriving at the NMJ during the refractory period of the muscle will fail to activate the muscle and are thus lost. To avoid losing spikes i.e. information and energy, instantaneous firing rates must stay below frequencies that would lead to lost spikes. Because the refractory period of male syrinx muscles ranges from 0.7 and 1.2 ms, motor neurons must fire below 1000 Hz. Together these two predictions set the lower and upper boundary for motor code spike rates. Lastly, we can further predict the peak of the distribution of nXIIts firing frequencies: Force increase by skeletal muscles is typically achieved in two ways: i) by increasing the number of recruited motor neurons and thus connected muscle fibers, and ii) by increasing the firing rate of motor neurons. These two mechanisms are known as recruitment and rate coding, respectively. Rate coding is most effective on the steep slope of the FFR (Fuglevand et al., 2015; Enoka and Duchateau, 2017). Additionally, to achieve unmodulated force increases, frequencies should be above *f_fused_*. Therefore, we predict ISIs between 3.4 ms (corresponding to an average *f_fused_* of 290 Hz) and 2.5 ms (average *f_Tet_*: 413 Hz) to be most suited to change force output of syrinx muscles in adult males. However, further experiments are needed to confirm that syringeal muscles and their innervating motor neurons are speed matched.

The hypothesis that speed matching occurs during the process of vocal development predicts that the ISI distribution of motor neurons in juveniles should be shifted towards higher ISIs to match the slower speed of the juvenile syringeal muscles. Due to the lack of data on motor neuron-firing we cannot test this hypothesis directly. However, the firing characteristics of the cortical premotor neurons projecting to the motor neurons in nXIIts (namely RA_nXIIts_) have been characterized over development (Olveczky et al., 2011). Indeed, the refractory period, *f_Tet_, f_fused_* and the frequency when force responses start to sum before and after song learning align well to the ISI probability distributions of age matched RA_nXIIts_ neurons during singing (Fig 5. *A, B*). The ISI distribution of spikes in RA becomes narrower from subsong (Fig. 5*A*) to crystallized adult song (Fig. 5*B*) and the boundaries are perfectly demarcated by t_Ref_ and frequency when force responses start to sum. The majority of pre-motor neuron ISIs lies between t_Ref_ and *f_fused_* for both age groups. Taken together, RA activity supports the notion that speed matching occurs over development.

**Figure 5:**
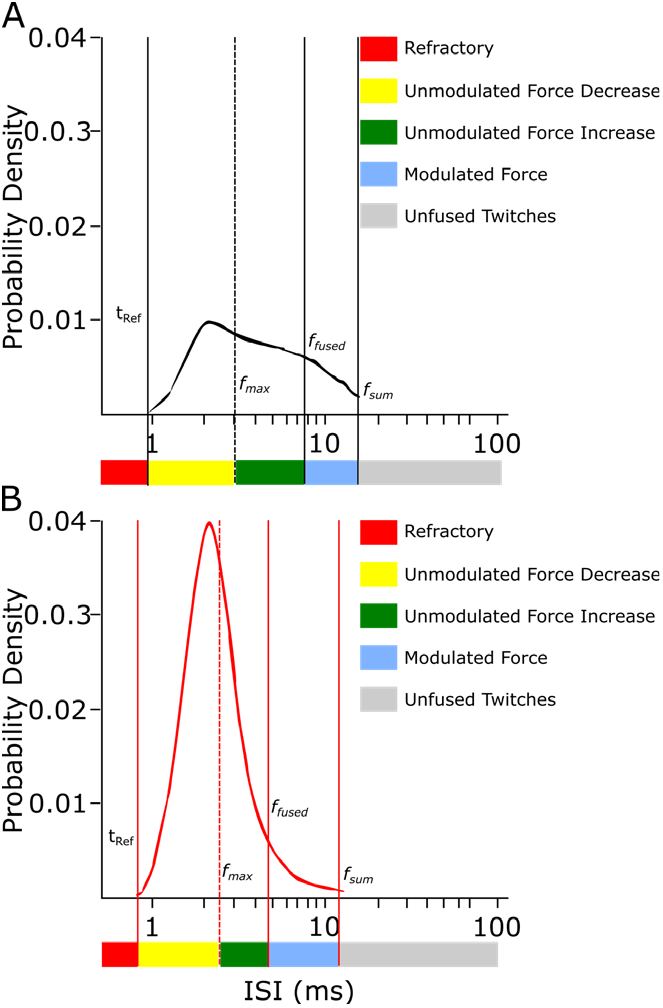
Biomechanical properties of syrinx muscles match the ISI distribution of the last premotor neurons driving them. The refractory period (t_Ref_) and summation threshold (*f_sum_*) of syrinx muscles match the boundaries of the RA-ISI distribution before the start of song learning (*A*) as well as after crystallization (*B*). The majority of ISIs lies between t_Ref_ and *f_fused_*. The ISI probability density plots are modified from (Olveczky et al., 2011).

Interestingly the refractory period of syrinx muscles is with <1 ms much shorter than the t_Ref_ of normal skeletal muscles and within the range of refractory times typically found in motor neurons (1-2 ms (Boerio et al., 2005)). We show that syringeal muscle is able to support instantaneous firing rates up to 1000 Hz, while normal skeletal muscles are typically driven by <50 Hz (Enoka and Duchateau, 2017). The only muscle activation rates we are aware of that are higher are those found in the electric organ of the weakly electric Brown ghost knife fish (*Apteronotus leptorhynchus*). The electrical organ contains heavily modified myofibers that lost the capacity to contract, and spinal neurons innervating them can reach brief bursts >1100 Hz (Dunlap et al., 2010). Thus, in addition to speed-limiting fundamental architectural constraints of skeletal muscle (Mead et al., 2017), extreme speed in songbird syrinx muscles may additionally be limited by another hard constraint: the refractory time of the central nervous system.

Our results support the idea that it is critical to include body biomechanics when interpreting motor commands (Nishikawa et al., 2007; Tytell et al., 2011) and emphasize the need for an embodied view of song motor learning (Düring and Elemans, 2016; Zhang and Ghazanfar, 2018). The increased higher variability of juvenile vocalizations (Tchernichovski et al., 2001; Olveczky et al., 2011) could for example be partially explained by a mismatch between contraction speed and neural firing characteristics during the process of vocal development (Adam and Elemans, 2019; Kollmorgen et al., 2020). Which mechanisms drive muscle speed changes over developmental timescales remains unknown, but we recently argued it may be driven predominantly by use, and not hormones (Adam and Elemans, 2019). This implies that extensive training of syringeal muscles is required to achieve maximal speed, and that the total duration and trajectory of song learning may thus not solely be set by neural circuit formation alone but may also be required for extensive muscle training.

## Acknowledgements

We would like to thank Sam Sober for comments on an earlier version of our manuscript. This work was supported by Danish Research Council (DFF 5051-00195), Carlsberg Foundation (CF17-0949) to I.A. and National Institute of Health (5R01NS099375-02) and NovoNordisk Foundation (NNF17OC0028928) to C.P.H.E

## References

Adam I, Elemans CPH (2019) Vocal Motor Performance in Birdsong Requires Brain-Body Interaction. eNeuro 6.

Adam I, Maxwell A, Rössler H, Hansen EB, Vellema M, Elemans CPH (2020) One-to-one innervation of vocal muscles allows precise control of birdsong. bioRxiv.

Binder MD, Heckman C, Powers RKJCP (2010) The Physiological Control of Motoneuron Activity. 3–53.

Boerio D, Hogrel JY, Creange A, Lefaucheur JP (2005) A reappraisal of various methods for measuring motor nerve refractory period in humans. Clin Neurophysiol 116:969–976.

Brainard MS, Doupe AJ (2013) Translating birdsong: songbirds as a model for basic and applied medical research. Annual review of neuroscience 36:489–517.

Chi Z, Margoliash D (2001) Temporal precision and temporal drift in brain and behavior of zebra finch song. Neuron 32:899–910.

Chiel HJ, Beer RD (1997) The brain has a body: adaptive behavior emerges from interactions of nervous system, body and environment. Trends Neurosci 20:553–557.

Dunlap KD, DiBenedictis BT, Banever SR (2010) Chirping response of weakly electric knife fish (Apteronotus leptorhynchus) to low-frequency electric signals and to heterospecific electric fish. J Exp Biol 213:2234–2242.

Düring DN, Elemans CPH (2016) Embodied Motor Control of Avian Vocal Production. In: Vertebrate Sound Production and Acoustic Communication (Suthers RA, Fitch WT, Fay RR, Popper AN, eds), pp 119–157. Cham: Springer International Publishing.

Elemans CP (2014) The singer and the song: the neuromechanics of avian sound production. Curr Opin Neurobiol 28:172–178.

Elemans CP, Mead AF, Rome LC, Goller F (2008) Superfast vocal muscles control song production in songbirds. PLoS One 3:e2581.

Elemans CP, Spierts IL, Muller UK, Van Leeuwen JL, Goller F (2004) Bird song: superfast muscles control dove’s trill. Nature 431:146.

Elemans CP, Rasmussen JH, Herbst CT, During DN, Zollinger SA, Brumm H, Srivastava K, Svane N, Ding M, Larsen ON, Sober SJ, Svec JG (2015) Universal mechanisms of sound production and control in birds and mammals. Nat Commun 6:8978.

Enoka RM, Duchateau J (2017) Rate Coding and the Control of Muscle Force. Cold Spring Harb Perspect Med 7.

Farmer TW, Buchthal F, Rosenfalck P (1960) Refractory period of human muscle after the passage of a propagated action potential. Electroencephalogr Clin Neurophysiol 12:455–466.

Fee MS, Scharff C (2010) The songbird as a model for the generation and learning of complex sequential behaviors. Ilar J 51:362–377.

Fitch WT, Huber L, Bugnyar T (2010) Social cognition and the evolution of language: constructing cognitive phylogenies. Neuron 65:795–814.

Fuglevand AJ, Lester RA, Johns RK (2015) Distinguishing intrinsic from extrinsic factors underlying firing rate saturation in human motor units. J Neurophysiol 113:1310–1322.

Goller F, Suthers RA (1996) Role of syringeal muscles in controlling the phonology of bird song. J Neurophysiol 76:287–300.

Hahnloser RH, Kozhevnikov AA, Fee MS (2002) An ultra-sparse code underlies the generation of neural sequences in a songbird. Nature 419:65–70.

Immelmann K (1984) The natural history of bird learning. Biol Learning, ed Berlin ea:271–288.

Jarvis ED (2004) Learned birdsong and the neurobiology of human language. Ann N Y Acad Sci 1016:749–777.

Kernell D (1995) Neuromuscular frequency-coding and fatigue. Adv Exp Med Biol 384:135–145.

Kollmorgen S, Hahnloser RHR, Mante V (2020) Nearest neighbours reveal fast and slow components of motor learning. Nature.

Leonardo A, Fee MS (2005) Ensemble coding of vocal control in birdsong. J Neurosci 25:652–661.

Lisberger SG, Medina JF (2015) How and why neural and motor variation are related. Curr Opin Neurobiol 33:110–116.

Lohr B, Dooling RJ (1998) Detection of changes in timbre and harmonicity in complex sounds by zebra finches (Taeniopygia guttata) and budgerigars (Melopsittacus undulatus). J Comp Psychol 112:36–47.

Manuel M, Chardon M, Tysseling V, Heckman CJ (2019) Scaling of Motor Output, From Mouse to Humans. Physiology (Bethesda) 34:5–13.

Mead AF, Osinalde N, Ortenblad N, Nielsen J, Brewer J, Vellema M, Adam I, Scharff C, Song Y, Frandsen U, Blagoev B, Kratchmarova I, Elemans CP (2017) Fundamental constraints in synchronous muscle limit superfast motor control in vertebrates. Elife 6.

Mendez J, Keys A (1960) Density and Composition of Mammalian Muscle. Metabolism 9:184–188.

Mooney R (2009) Neural mechanisms for learned birdsong. Learn Mem 16:655–669.

Nishikawa K, Biewener AA, Aerts P, Ahn AN, Chiel HJ, Daley MA, Daniel TL, Full RJ, Hale ME, Hedrick TL, Lappin AK, Nichols TR, Quinn RD, Satterlie RA, Szymik B (2007) Neuromechanics: an integrative approach for understanding motor control. Integr Comp Biol 47:16–54.

Nottebohm F, Stokes TM, Leonard CM (1976) Central control of song in the canary, Serinus canarius. J Comp Neurol 165:457–486.

Olveczky BP, Otchy TM, Goldberg JH, Aronov D, Fee MS (2011) Changes in the neural control of a complex motor sequence during learning. J Neurophysiol 106:386–397.

Orizio C, Gobbo M, Diemont B (2004) Changes of the force-frequency relationship in human tibialis anterior at fatigue. J Electromyogr Kinesiol 14:523–530.

Otchy TM, Michas C, Lee B, Gopalan K, Gleick J, Semu D, Darkwa L, Holinski BJ, Chew DJ, White AE, Gardner TJ (2019) Printable microscale interfaces for long-term peripheral nerve mapping and precision control. bioRxiv.

Raikova RT, Aladjov H (2002) Hierarchical genetic algorithm versus static optimization-investigation of elbow flexion and extension movements. J Biomech 35:1123–1135.

Riede T, Goller F (2010) Peripheral mechanisms for vocal production in birds - differences and similarities to human speech and singing. Brain Lang 115:69–80.

Ritz C, Baty F, Streibig JC, Gerhard D (2015) Dose-Response Analysis Using R. PLoS One 10:e0146021.

Roper A, Zann R (2006) The onset of song learning and song tutor selection in fledgling zebra finches. Ethology 112:458–470.

Scharff C, Nottebohm F (1991) A comparative study of the behavioral deficits following lesions of various parts of the zebra finch song system: implications for vocal learning. J Neurosci 11:2896–2913.

Signorell A, al. m (2019) DescTools: Tools for Descriptive Statistics. R Foundation for Statistical Computing, Vienna, Austria.

Sober SJ, Wohlgemuth MJ, Brainard MS (2008) Central contributions to acoustic variation in birdsong. J Neurosci 28:10370–10379.

Sober SJ, Sponberg S, Nemenman I, Ting LH (2018) Millisecond Spike Timing Codes for Motor Control. Trends Neurosci 41:644–648.

Srivastava KH, Elemans CP, Sober SJ (2015) Multifunctional and Context-Dependent Control of Vocal Acoustics by Individual Muscles. J Neurosci 35:14183–14194.

Srivastava KH, Holmes CM, Vellema M, Pack AR, Elemans CPH, Nemenman I, Sober SJ (2017) Motor control by precisely timed spike patterns. P Natl Acad Sci USA 114:1171–1176.

Stein RB, Gossen ER, Jones KE (2005) Neuronal variability: noise or part of the signal? Nat Rev Neurosci 6:389–397.

Tang C, Chehayeb D, Srivastava K, Nemenman I, Sober SJ (2014) Millisecond-scale motor encoding in a cortical vocal area. PLoS Biol 12:e1002018.

Tchernichovski O, Mitra PP, Lints T, Nottebohm F (2001) Dynamics of the vocal imitation process: how a zebra finch learns its song. Science 291:2564–2569.

Tytell ED, Holmes P, Cohen AH (2011) Spikes alone do not behavior make: why neuroscience needs biomechanics. Curr Opin Neurobiol 21:816–822.

Watanabe S, Kitawaki T, Oka H (2010) Mathematical equation of fusion index of tetanic contraction of skeletal muscles. J Electromyogr Kinesiol 20:284–289.

Wild JM (1993a) Descending projections of the songbird nucleus robustus archistriatalis. J Comp Neurol 338:225–241.

Wild JM (1993b) The avian nucleus retroambigualis: a nucleus for breathing, singing and calling. Brain Res 606:319–324.

Wild JM (1997) Neural pathways for the control of birdsong production. J Neurobiol 33:653–670.

Williams H, Nottebohm F (1985) Auditory responses in avian vocal motor neurons: a motor theory for song perception in birds. Science 229:279–282.

Zhang YS, Ghazanfar AA (2018) Vocal development through morphological computation. PLoS Biol 16:e2003933.

Zhang YS, Takahashi DY, Liao DA, Ghazanfar AA, Elemans CPH (2019) Vocal state change through laryngeal development. Nat Commun 10:4592.

Zhurov Y, Brezina V (2006) Variability of motor neuron spike timing maintains and shapes contractions of the accessory radula closer muscle of Aplysia. J Neurosci 26:7056–7070.

